# A Quantitative Hematopoietic Stem Cell Reconstitution Protocol: Accounting for Recipient Variability, Tissue Distribution and Cell Half-Lives

**DOI:** 10.1101/2020.12.03.410894

**Authors:** Smrithi Rajendiran, Scott W. Boyer, E. Camilla Forsberg

## Abstract

Hematopoietic stem and progenitor cell (HSPC) transplantation is the paradigm for stem cell therapies. The protocol described here enables quantitative assessment of the body-wide HSPC reconstitution of different mature hematopoietic cells in mice based on their presence in circulating blood. The method determines donor-derived mature cell populations per mouse, over time, by quantitatively obtaining their absolute numbers in the peripheral blood and utilizing previously assessed tissue-distribution factors. A Markov-based birth/death computational model accounts for the drastic differences in mature cell half-lives. By quantifying the number of cells produced and eliminating host variability, the protocol can be used to directly compare the lineage output of different types of HSPCs on a per cell basis, thereby clarifying the lineage potential and expansion capacity of different cell populations. These protocols were developed for hematopoiesis, but can readily be extended to other contexts by simply replacing the cell types and distributions.

**Highlights:**

- Quantitative assessment of stem and progenitor cell reconstitution capacity
- Elimination of cell-specific recipient variability for accurate donor cell potential
- Directly comparable lineage output within and between stem and progenitor cells
- Blood-based absolute quantification of whole-body repopulation over time
- Markov modelling-based consideration of differential mature cell half-lives

## Introduction

Transplanting cells from one entity, the donor, to another, the recipient, has been attempted since the early 1900s. In 1956, Dr. E. Donnall Thomas and colleagues successfully engrafted hematopoietic cells into a leukemic patient from the patient’s identical twin. The field of hematopoietic transplantation has since progressed such that it is now possible to transplant cells from non-identical twins and even unrelated individuals (Gyurkocza and Sandmaier, 2014; Jenq and Van Den Brink, 2010; Juric et al., 2016; Singh and McGuirk, 2016).

Much of the progress towards clinical success is owed to mechanistic studies in non-human organisms. Mouse models are widely used in the study of hematopoiesis because the underlying mechanisms closely mirror those of human hematopoiesis (Boieri et al., 2016; Sykes and Scadden, 2013). High conservation is observed in the types of HSPC populations present, the organ sites of hematopoietic cell generation and maturation, and the regulation of HSPC trafficking between organs within an individual and upon transplantation. Hematopoietic cell transplantations are potentially life-long, curative treatments for a broad range of blood disorders (Bair et al., 2020; Smith-Berdan et al., 2019; Strocchio and Locatelli, 2018; Tanhehco and Bhatia, 2019). Their success depends on robust and prolonged generation of blood and immune cells. Consequently, tremendous efforts have been dedicated to understand the lineage potential and expansion capacity of the cells that comprise a transplant. Despite decades of work, significant controversies persist with regards to lineage potential and bias, and the relationship between different progenitor populations (Boyer et al., 2011; Kondo, 2010; Perié et al., 2015; Woolthuis and Park, 2016). These issues have been exacerbated by the difficulty in directly tracking donor-derived mature red blood cells (RBCs) and platelets (Plts). In an effort to resolve these longstanding issues, we have developed protocols for improved quantitative assessment of HSPC-derived cells, including RBCs, Plts, granulocytes-macrophages (GM), B and T cells.

Historically, assessment of HSPC reconstitution capacity in experimental models have mainly focused on donor chimerism (Bader et al., 2005; Khan et al., 2004): the proportion of cells that are donor-derived versus those that are produced by the host HSPCs. A major drawback of chimerism data is the tremendous variation in *host* cell numbers due to the requisite preconditioning of the recipient: we recently reported that recipient B cells were rapidly reduced by ~2,000-fold upon lethal preconditioning, whereas RBCs underwent a slower and drastically lower (~4-fold) decline (Boyer et al., 2019). Thus, “donor chimerism” is heavily dependent on the cell type-specific and time- and dose-dependent reduction and recovery of recipient hematopoiesis. This results in a skewed perception of the lineage potential of the transplanted cell population (Boyer et al., 2019). To enable a more accurate view of the cell production capacity of transplanted cells, we developed a quantitative protocol that eliminates the host cell variable (Rajendiran et al., 2020). We also assessed the tissue distribution of each mature cell type, so that measurements made by peripheral blood sampling can be extrapolated to whole-body reconstitution. Additionally, we implemented birth/death Markov modeling to account for the drastically different half-lives of mature hematopoietic cell types (ranging from ~1 day to several months) to decouple cell accumulation from active cell production (E. and Yule, 1925; Kendall, 1948). This new method makes it possible to quantitatively track all HSPC-produced cells in a recipient over time based solely on blood sampling.

### Development of the method

To better understand HSPC capacity to produce the 5 major terminally differentiated cell types within the 4 broad lineages, i.e., erythroid (Red Blood Cells), megakaryocytic (Platelets), myeloid (Granulocyte Myelomonocyte/Macrophages), and lymphoid (B-cells and T-cells), we established a new method that enables quantification of absolute number of donor-derived cells based on their numbers in the peripheral blood and the distribution within the entire recipient. We quantified the number of terminally differentiated cells, including RBCs and Platelets, in the major hematopoietic compartments (blood, bone marrow, spleen, lymph nodes and thymus) and then utilized this tissue distribution to infer the total numbers of each of the mature cells of donor origin in the recipient mouse post-transplantation of various HSPC populations (HSCs, MPPs, CMPs, GMPs, MEPs and CLPs) based on blood data alone (Boyer et al., 2019). Using Markov modeling, this protocol also accounts for differences between the number of cells generated and cells accumulated over time based on their varying half-lives. Together, these calculations enable direct, quantitative cross-comparison of the per-cell potential of distinct HSPCs.

### Assumptions and Limitations

We analyzed 5 hematopoietic tissues, namely blood, bone marrow (BM), spleen, lymph nodes (LN) and thymus as the primary tissues that contain the mature cells (i.e., RBC, Plt, GM, B and T cells). These are known to contain the majority of mature hematopoietic cells, but some mature cells could also reside in other tissues, such as platelets in the lungs (Lefrançais et al., 2017), which is not accounted for in this study. The assumptions reported here and used previously (Boyer et al., 2019), were based on data from a mix of young, adult male and female mice that are about 8-12 weeks of age (and weighing ~20g-25g) at the time of using them for transplants. If either the recipient or donor mouse is of a different strain, age, sex and/or genetically modified, the cell distribution may be different than what we report for wild-type C57BL/6 mice; this could be tested by methods analogous to those described here and previously (Boyer et al., 2019). Additionally, the assumption that the BM from 2 legs (femurs plus tibias) of the mouse accounts for 25% of the total bone marrow and that the three lymph node pairs that we extracted here accounts for about 20% of all the lymph nodes are approximations based on literature (Dunn, 1954). If more specific numbers are desired, the protocol can be adjusted to include all the bones for BM and all the lymph nodes for the total mature cells in these compartments. We also presumed that the donor-derived cells upon transplantation assume the same tissue distribution as the cells in untreated mice.

To model the cell generation rate for the 5 mature cells, we used literature-based half-lives (Dholakia et al., 2015; Fulcher and Basten, 1997; Nayak et al., 2013; Simon and Kim, 2010; Sprent and Basten, 1973; Swirski et al., 2006; Wen et al., 2013), and we made the assumption that cell half-lives are similar upon transplantation. More accurate assessments could be made if half-lives were tested for each specific condition. The model can readily be extended to other cell types with known half-lives with minimal modification to the current script. The overall protocol can be extended to include other tissues and/or cell types if the experimental procedures include those cell types/tissues.

### Applications of the method

Here, we have provided a means to obtain absolute quantification in the 5 main hematopoietic tissue compartments for the 5 major mature cell types from different donor cells post-transplantation instead of relying on donor cell contribution in terms of chimerism alone. This method revealed new insights about lineage potential of various hematopoietic stem and progenitor cells (HSPCs) by enabling direct, quantitative comparison of cell reconstitution capacity (Boyer et al., 2019). Using this method, we can also track the changes in the absolute numbers of mature cells present over time. In addition, we can make an approximate assessment of the cellular output per transplanted cell. Two illustrative examples from that study are reproduced here (**Figures 1 and 2**). Figure 1 shows how the donor-derived mature cell contributions by transplanted MPP^F^s as traditional chimerism (Figure 1A) appear drastically different when transformed into absolute numbers of donor-derived cells per microliter of peripheral blood (Figure 1B) or in the entire mouse per transplanted input cell (Figure 1C). **Figure 2** demonstrates that these different views depend on both the host- and donor-derived mature cells. The magnitude of host B-cell depletion (Figure 2A) upon preconditioning irradiation is almost 10-fold greater than that of host T-cells (Figure 2B). This effect is compounded by the more rapid production of B-cells (Figure 2A) compared to T-cells (Figure 2B) from transplanted CLPs. (Boyer et al., 2019). Implementation of the Markov modelling that accounts for cell half-life allows discrimination between cells accumulated versus cells generated over time, as demonstrated in Boyer et al. 2019 (Boyer et al., 2019). **Figure 3** shows that the mature cell half-lives do not significantly alter the perceived reconstitution capacity of transplanted cells (Figure 3B), because the majority of cells are “born” (produced) very soon after transplantation (Figure 3A).

**Figure 1:**
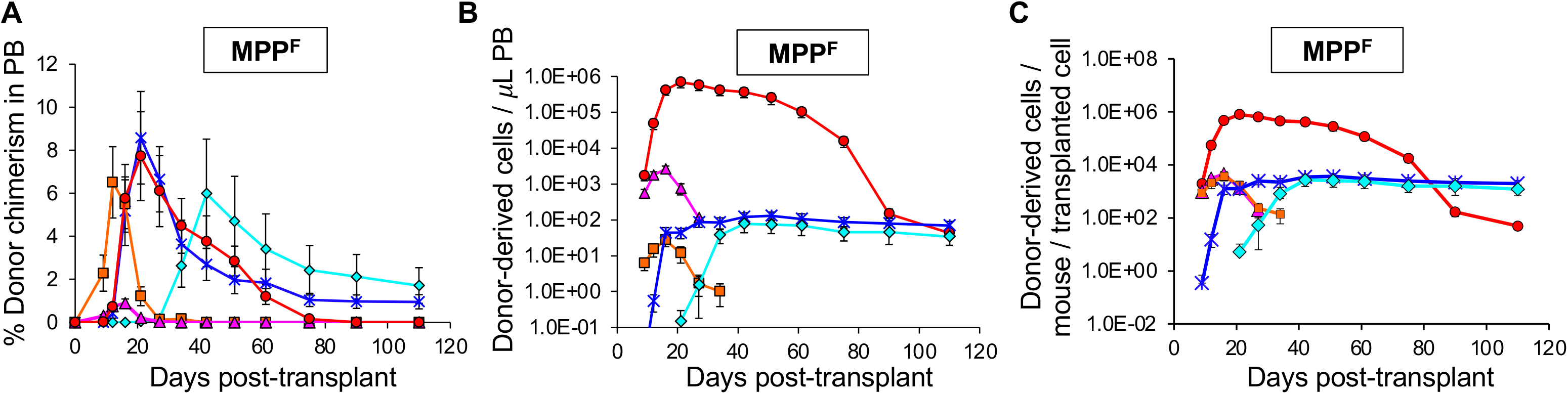
Converting chimerism data to absolute quantification drastically alters the perceived reconstitution potential of transplanted stem and progenitor cells. Visualization of the differences between donor chimerism (A), and absolute donor-derived cells in the peripheral blood (B) and in the entire recipient (C). Reconstitution potential of multipotent progenitor cells (1,000 MPP^F^/recipient; MPP^F^ were isolated as ckit++lineage-Sca1++CD150-Flk2+ bone marrow cells from UBC-GFP mice) upon transplantation into sublethally irradiated (500 rad) wild-type mice. **(A)** Percent donor chimerism in the peripheral blood (PB) over 110 days from 1,000 MPP^F^. **(B)** Reconstitution data from (**A**) replotted as the absolute number of donor-derived cells per microliter PB. **(C)** Reconstitution data from (**A and B**) replotted as the absolute number of donor-derived mature cells per transplanted MPP^F^ in the entire mouse. Data are means ± SEM from at least seven recipients and two independent experiments. Results are examples from Boyer et al, 2019.

**Figure 2:**
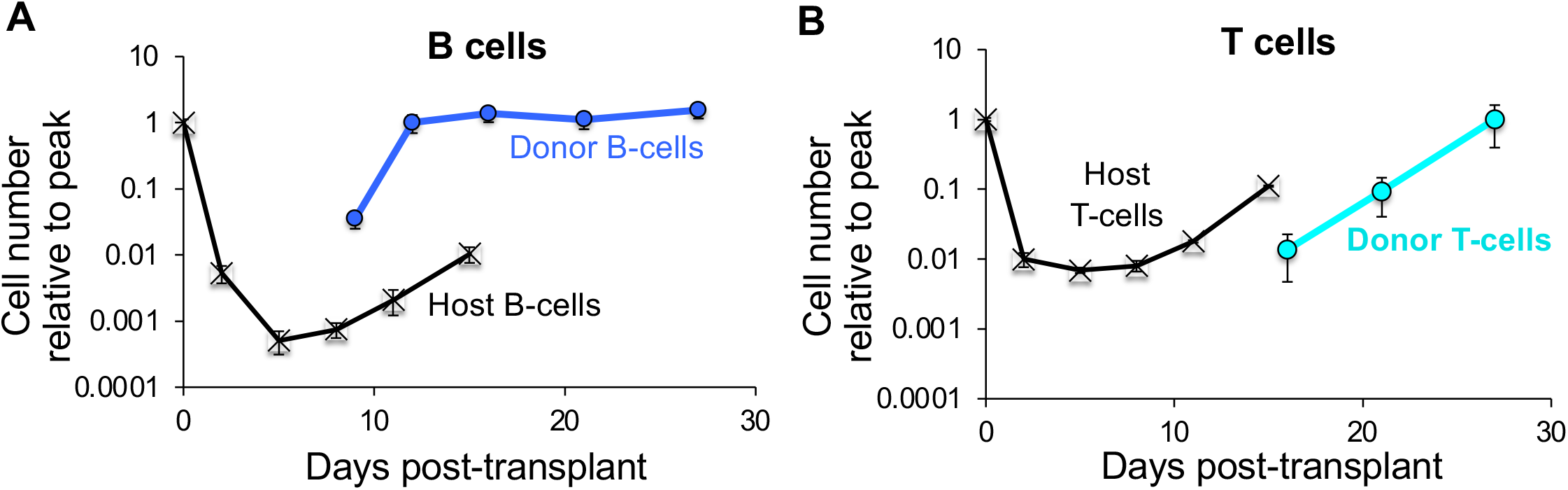
CLP^F^-derived B cells accumulate near the low point of host B cell decline, whereas host T cells recover prior to CLP-derived T cell accumulation. Black lines depict the decline and recovery of host B cells (**A**) and T cells (**B**) after lethal irradiation. Blue and green lines indicate donor-derived B cells (**A**) and T cells (**bottom**), respectively, after transplantation of 10,000 CLP^F^ (isolated as lineage-IL7R+Flk2+ckit+Sca1+ bone marrow cells from UBC-GFP mice) into sublethally irradiated (500 rad) wild-type mice. Data are means ± SEM from at least seven recipients and two independent experiments. Results are examples from Boyer et al, 2019.

**Figure 3:**
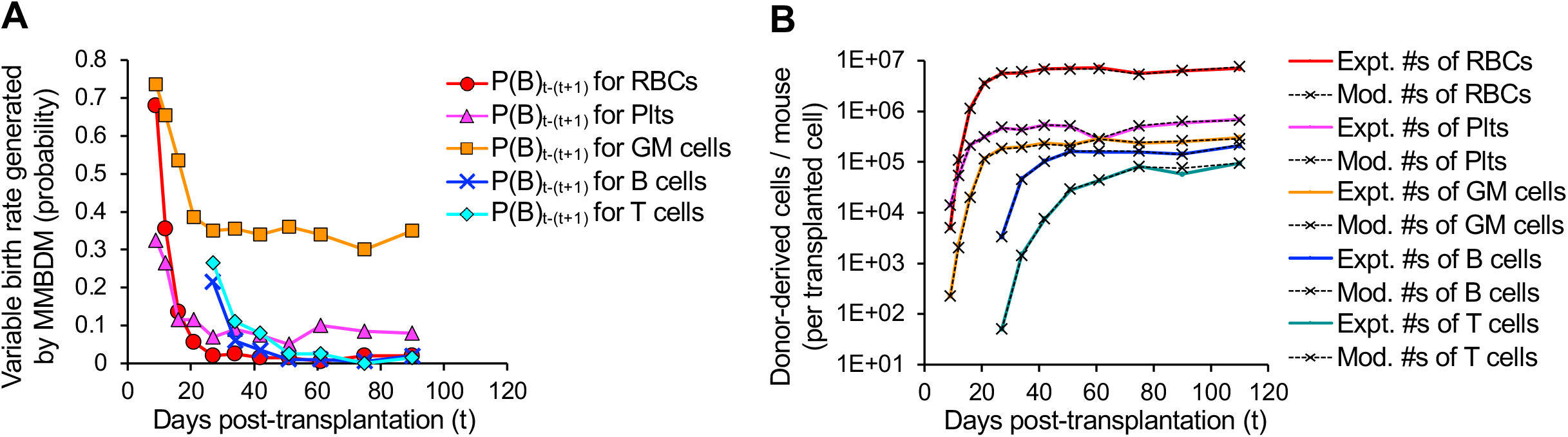
Markov-based modeling of birth-death probabilities and the effects on reconstitution profiles. Example of data generated by Modified Markov Birth-Death Model (MMBDM) for both the estimated birth probabilities (**A**) and the corresponding model fitted cell number values (**B**). **(A)** The MMBDM was used to estimate the birth rate (probabilities) between two experimentally assessed time points by taking into account the assumed death rates (probabilities) from literature for each of the 5 mature lineages and the number of cells detected experimentally at both the time points. The birth probability (plotted here) is calculated depending on the closest fit of the model generated numbers to the actual number of cells present at the two time points. The birth rate (in days) can then be estimated between successive time points as log 2 base e divided by birth probabilities generated. P(B)_t–(t+1)_= Birth rate probability plotted at time t, for time between t and t+1. **(B)** Experimentally observed donor cell derived mature cells per mouse compared to the numbers generated by MMBDM. The overlapping curves (colored [experimental] line with the corresponding black dashed [modeled] line for each population) shows that we achieved high accuracy using the MMBDM to estimate the number of cells present at the various time points. Expt. #s = Experimentally derived numbers at time t; Mod. #s = MMBDM-based numbers at time t.

We first used this absolute quantification method with respect to RBC, Plt, GM, B and T cells in the hematopoietic system (Boyer et al., 2019), but analogous techniques can be applied to other tissues and cell types (Cool et al., 2020). The method can also be used to compare the competitive fitness between wild-type and mutated cells. Upon transplantation of other types of stem or progenitor cells, this method can be used to track the absolute *in vivo* potential of the transplanted cell and the donor-derived differentiated progeny cell numbers. This method will provide new insights about lineage bias and donor cell potential independent of host related factors.

### Reagents/Instruments/Software

Mice (wild-type C57BL/6 and UBC-GFP donor mice), scissors, scalpels, forceps, Eppendorf tubes, pipette, pipette tips, FACS tubes (polypropylene or polystyrene), Kimwipes, APC Calibrite Beads, 20mM EDTA in 1X PBS, donor calf serum, 5mM EDTA in 1X PBS with 2% donor calf serum (staining media, SM), ACK lysis buffer, blocking agent (rat IgG), anti-mouse antibodies, cell viability dye like propidium iodide (PI), mortar and pestle, dounce homogenizer, 70μm nylon mesh filters (filters), CO_2_ chamber to sacrifice mice, isoflurane to anesthetize during retro-orbital injections, 23 and 27½ G needles, timer, Flow Cytometer (with ability to detect 8 or more fluorophores), irradiator (or Busulfan); Fluorescence Activated Cell Sorter (for sorting if transplanting purified cells, and analysis of the mature cells); flow cytometry acquisition and analysis software such as Diva/FlowJo; Microsoft Excel; Python.

### Protocol/Procedure

#### Protocol for measurement of RBCs, Platelets, GM cells, B cells and T cells in various tissues

~ 75 minutes

The goal of this step is to assess the distribution of the various mature cells in the different tissues and conversion to total number of cells in the mouse

1. C57BL/6 mice, 8-12 weeks of age and 20-25g by weight, were sacrificed by CO_2_.
2. Mature cells from different tissues were collected:
  a. Peripheral/circulating blood (total blood collected from a mouse by perfusion): ~ 5 minutes
    aa. Prepare a syringe with 15-20mL of 20mM EDTA in PBS and a 23G needle.
    ab. Prop the mouse on its back and pin it down by its arms with needles.
    ac. Spray the skin (to prevent fur interference and keep the area sterile) with ethanol and cut above the diaphragm. Be careful to not sever any blood vessels.
    ad. Hold the sternum, make a shallow cut just below it (diaphragm needs to be intact to collect the blood) and cut beside it, on either side, towards the face of the mouse. Also, make sure to hold this mid piece in place with a pin/needle to allow unrestricted access to the heart in the chest cavity between the diaphragm and the neck.
    ae. To prevent clotting, add 300μL 20mM EDTA to the cavity. Cut the right atrium of the heart to let the blood flow into the cavity.
    af. Inject the remaining 20mM EDTA, slowly over about 2 to 3 minutes, through the right ventricle and use a pipet to collect the blood that accumulates into the chest cavity.
    ag. By the end of the perfusion, the blood that comes out should be much paler (clearer) than in the beginning since the 20mM EDTA replaces the blood in the vessels. Place the tube with collected perfused blood on ice.
  b. Bone marrow from the hind limbs (femurs and tibia) two long bones in the legs: ~ 5 minutes
    ba. Spray the skin around the thighs and legs with ethanol, cut the skin and cut the muscle along the femur, towards the hip joint.
    bb. Place the scissor under the patella and snap the femur from the hip at the joint. Cut along the bone (do not cut bone) to remove as much fat as possible and place the entire leg in SM on ice. Repeat with the femur on the other side.
    bc. Clean the fat/muscle from the bones with forceps/scissors/Kimwipes and place back in fresh SM.
    bd. Crush the bones (gently, and till the crunching sound can no longer be heard) with a mortar and pestle, in SM. Gently pipet up and down to dislodge the bone marrow cells. Transfer the solution through a filter into a FACS tube. Repeat crushing and cleaning the bones to get all the cells until the SM after all the washes is clear and the bones are almost white.
    be. Take care not to over-crush the bones (this can affect cell viability). Place the filtered solution in ice.
  c. Spleen: ~ 5 minutes
    ca. Spray ethanol on the side of the mouse and cut the spleen out. Place it in SM on ice.
    cb. Crush spleen using a dounce homogenizer or mortar and pestle with SM. Pass cells through a filter into a FACS tube on ice.
  d. Lymph nodes: ~ 5 minutes
    da. Remove the superficial cervical, inguinal and axillary lymph nodes.
    db. Crush the lymph nodes in SM using a dounce homogenizer or mortar and pestle and pass the sample through a filter into a FACS tube on ice.
  e. Thymus: ~ 5 minutes
    ea. Remove the thymus.
    eb. Crush the thymus in SM using a dounce homogenizer or mortar and pestle and pass through a filter into a FACS tube on ice.
3. Add preset/pre-recorded number of APC Calibrate Beads to each of the 5 samples such that at least 1,000 beads can be collected per million cells. Spin down the cells from the different tissues at 1200rpm for 5 minutes at 4°C.
4. Remove the supernatant carefully by aspiration or with a pipette. Re-suspend pellets in SM to make a single-cell suspension. These samples will be used to calculate the total number of the various mature cells in the respective tissues.
5. The assumptions about the fractions of tissues collected by the above preparations are shown in **BOX 1a**.
6. Transfer a fraction of each of the samples to new FACS tube labeled with the respective tissue names, after ensuring that the samples are mixed well with APC beads. Block with rat IgG in SM for 10 minutes on ice. Stain with Ter119-Fl1, CD61-Fl2, Mac1-Fl3, Gr1-Fl4, B220-Fl5, CD3-Fl6 (Fl = fluorophore conjugate) for 20 minutes on ice in the dark. Wash with SM and spin down at 1200rpm for 5 minutes at 4°C. Also stain appropriate compensation controls.
7. Analyze the cells using a flow cytometer: RBCs, Platelets, GM cells, B-cells and T-cells.
8. The number of beads collected with the various mature cells in each sample can be detected in the APC channel in the flow cytometer.
9. The different mature cells in the individual tissues can be calculated using the following equation and represented as shown in **BOX 1b**, using BOX 1 assumptions.

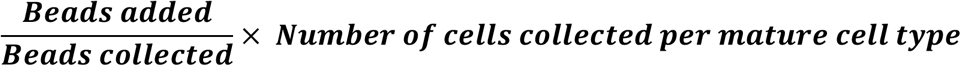
10. This provides the distribution of the different mature cells in the 5 tissues as shown in our publication in 2019 (Figure 1J and 1K of Boyer et al., 2019).
11. Next, we utilized numbers in the circulating blood and numbers in the total blood (for each cell type), along with the cells in the other tissues, to obtain a scaling factor for blood based total cell number calculation (fraction) as shown in **BOX 2**.

#### Protocol for Quantitative Tail Bleeds

~ 40 minutes

12. Prepare antibody/bead master mix (**BOX 3**) before bleeding the mice. Also prepare tubes for appropriate compensation controls.
13. Add 25μl of 20mM EDTA with 2% DCS and rat IgG to FACS tubes and keep covered on ice.
14. Place the mice that need to be bled into a cage with the heating lamp. Mark the tubes with the identifier while mice are warming up.
15. Once the mice are warm enough, nick the tail vein to collect blood in a 1.5ml Eppendorf tube and transfer 25μl of blood to the prepared FACS tube for the respective mouse. If less than 25μl is collected, note the volume on the FACS tube. Mix samples immediately and well to ensure no clots form. Incubate mixed blood samples for 10 minutes on ice.
16. After incubation, add 50μl of antibody/bead cocktail to samples, mix well and incubate for at least 20 minutes, protected from light, on ice. Ensure to also stain a sample that will allow donor/host distinction with the antibody/bead cocktail.
17. After incubation, add 1ml of SM to FACS tubes with the blood and mix well (without introducing any air bubbles) to ensure an even distribution of beads and cells.
18. Transfer 900μl to a new FACS tube labeled with the respective sample identifier (this is the fraction that needs to undergo ACK lysis). Add SM to both the sets of tubes to wash. Spin at 1200rpm at 4°C for 5 minutes and aspirate supernatant. Re-suspend whole blood fraction in 300μl of SM and leave in ice. This fraction is now ready to run on the cytometer.
19. Lyse the ACK fraction pellet in 1ml 1X ACK lysis buffer at room temperature, until samples become translucent.
20. Stop ACK reaction by adding SM (at least 3-4 times the volume of ACK used) and wash. Spin, discard the supernatant and resuspend in SM. This fraction is now ready to run on the cytometer.
21. On the flow cytometer: Whole Blood Sample – Record at least 1 million total events at a low FSC threshold to detect platelets and RBCs (for ex. 500 on our BD FACS Aria IIu or LSRII). ACK Lysed Sample – Record at least 1,000 GMs at a higher FSC threshold to detect nucleated cells (for ex. 10,000 on our BD FACS Aria IIu or LSRII).
22. Analyze as shown in **BOXES 4, 5 and 6** to obtain the traditional donor chimerism as well as the quantitative absolute number of donor-derived mature cells per μl of peripheral blood.

#### Protocol for total donor cell assessment in blood upon transplantation without sacrificing the mouse

23. After transplanting mice, perform bleeds like in Step 12 – 22 at desired time points post-transplantation.
24. Obtain the cells per μl (and the donor chimerism, if interested) as shown in **BOXES 4 – 6**.
25. Since repetitive blood measurements are required, it is not possible to sacrifice the mice to obtain the total number of mature cells at each time point. Hence, we derived a multiplication factor that can be used at each time point, using the distribution and scaling factors from **BOX 2** as shown in **BOX 7**. Multiplication of the absolute number of cells per μl from the tail bleed (R, P, G, B, T) as obtained in **BOX 6** can then be used in the formulae in **BOX 7** to provide the donor derived cells for each individual mature cell type per mouse.
26. Divide the numbers from step 25 by the total number of donor cells transplanted to obtain the average number of donor-derived mature cells per mouse per transplanted cell (Boyer et al 2019).

#### Protocol to distinguish between cells accumulated and generated based on the number of cells present at a given time

27. Markov Birth-Death Models (MBDM) have been used to allow estimation of changes in population size based on “birth” and “death” of the cells within the population. Here, for the 5 mature cells used, based on literature, assumptions about a constant half-life (death rate) have been made. Since the donor cells transplanted can potentially give rise to the mature cells (through multiple steps), and the mature cells can continue to live as is or die, we modified the Markov Birth-Death Model to allow dynamic changes to the birth rates of the mature cells form the donor cells over time resulting in a Modified Markov Birth-Death Model (MMBDM) as shown in **BOX 8**.
28. Based on the experimental data we obtained over time post transplantation, using the appropriately tuned Markov Birth-Death Model for each mature cell type of interest, we generated a program (https://github.com/cforsberg/Boyer-Stem-Cell-Reports-2019) that allows estimation of the variable birth rates and the number of “new cells generated” at the different time points for each of the 5 cell types. Given the close fit between the modeled curve and the experimental curve for the cells present and detected at any given time (**Figure 3**), this program is robust enough to allow reasonably accurate projections of the dynamic birth rates of the mature cell types post transplantation of HSPCs. Depending on the potency (differentiation potential) of the stem and progenitor cell in the hematopoietic hierarchy, persistent or short bursts of new cells are produced post transplantation, as demonstrated by Figure 2 in Boyer et al. 2019.

### Anticipated results/Concluding remarks

This novel quantitative method allows the absolute quantification of donor-derived terminally differentiated hematopoietic cells, independent of the host cell variability and/or pre-conditioning. It allows for direct comparison between different types of HSPCs’ ability to produce the 5 mature cell types *in vivo* (**Figures1–2**), considering the half-lives of the cells and the plasticity of the HSPCs (**Figure 3**) (Boyer et al., 2019). Given that the *in vivo* capabilities of the donor cells are directly influenced by multiple external factors in the host, the ability to get an unbiased and complete account of the total number of the terminally differentiated cells of all 5 lineages is important for accurately understanding the potential of the HSPCs and making comparisons between them.

This method can be extended to various disease scenarios that involve transplantation therapies, hematopoietic and non-hematopoietic, to allow better selection of the donor cells and appropriate pre-conditioning regimen(s) in the recipients. This understanding will eventually allow the recipients of stem cell transplantations to obtain the maximal benefit from the transplantation procedure.

## Author contributions

All authors read and approved the final manuscript.

Smrithi Rajendiran: Conceptualization, Methodology development, Writing, Reviewing and Editing

Scott Boyer: Conceptualization, Methodology development, Reviewing and Funding acquisition Camilla Forsberg: Supervision, Conceptualization, Methodology development, Writing, Reviewing, Editing and Funding acquisition

## Acknowledgements

We thank Praveen K. Muthuswamy for assistance with the Python code. We thank Forsberg lab members for comments on the manuscript. We also thank Bari Nazario at the UCSC Institute for the Biology of Stem Cells (IBSC) for flow cytometry support.

### BOX 1a: Absolute cell numbers from the different tissues

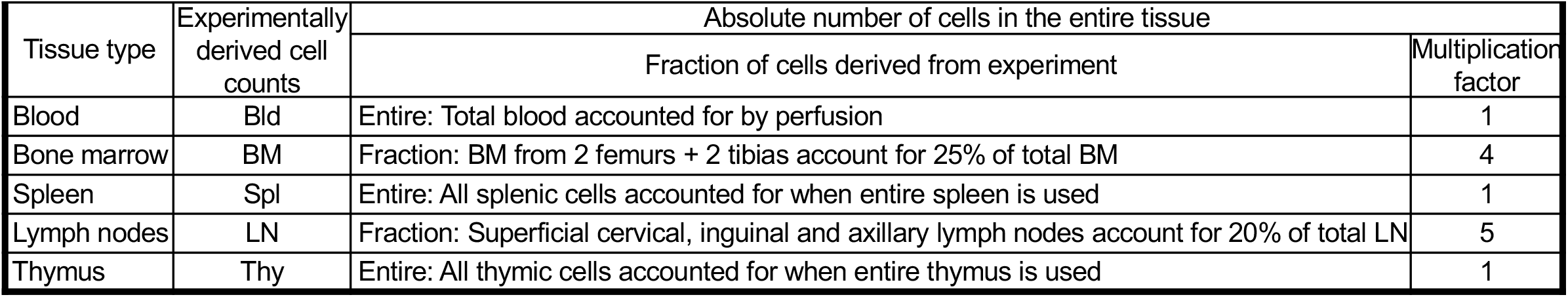

| BOX 1a: Absolute cell numbers from the different tissues |  |  |  |
| --- | --- | --- | --- |
| Tissue type | Experimentally derived cell counts | Absolute number of cells in the entire tissue |  |
|  |  | Fraction of cells derived from experiment | Multiplication factor |
| Blood | Bld | Entire: Total blood accounted for by perfusion | 1 |
| Bone marrow | BM | Fraction: BM from 2 femurs + 2 tibias account for 25% of total BM | 4 |
| Spleen | Spl | Entire: All splenic cells accounted for when entire spleen is used | 1 |
| Lymph nodes | LN | Fraction: Superficial cervical, inguinal and axillary lymph nodes account for 20% of total LN | 5 |
| Thymus | Thy | Entire: All thymic cells accounted for when entire thymus is used | 1 |

### BOX 1b Experimental

| BOX 1b | Experimental |  |  |  |  |
| --- | --- | --- | --- | --- | --- |
| Cells → | RBC | Plts | GM cells | B cells | T cells |
| Cells (Perfused blood) | $R_P$ | $P_P$ | $G_P$ | $B_P$ | $T_P$ |
| Cells (BM) | $R_B$ | $P_B$ | $G_B$ | $B_B$ | $T_B$ |
| Cells (Spleen) | $R_S$ | $P_S$ | $G_S$ | $B_S$ | $T_S$ |
| Cells (Lymph nodes) | $R_{LN}$ | $P_{LN}$ | $G_{LN}$ | $B_{LN}$ | $T_{LN}$ |
| Cells (Thymus) | $R_T$ | $P_T$ | $G_T$ | $B_T$ | $T_T$ |
| Total cells | $R_P + 4R_B + R_S + 5R_{LN} + R_T$ | $P_P + 4P_B + P_S + 5P_{LN} + P_T$ | $G_P + 4G_B + G_S + 5G_{LN} + G_T$ | $B_P + 4B_B + B_S + 5B_{LN} + B_T$ | $T_P + 4T_B + T_S + 5T_{LN} + T_T$ |

### BOX 2 Experimental

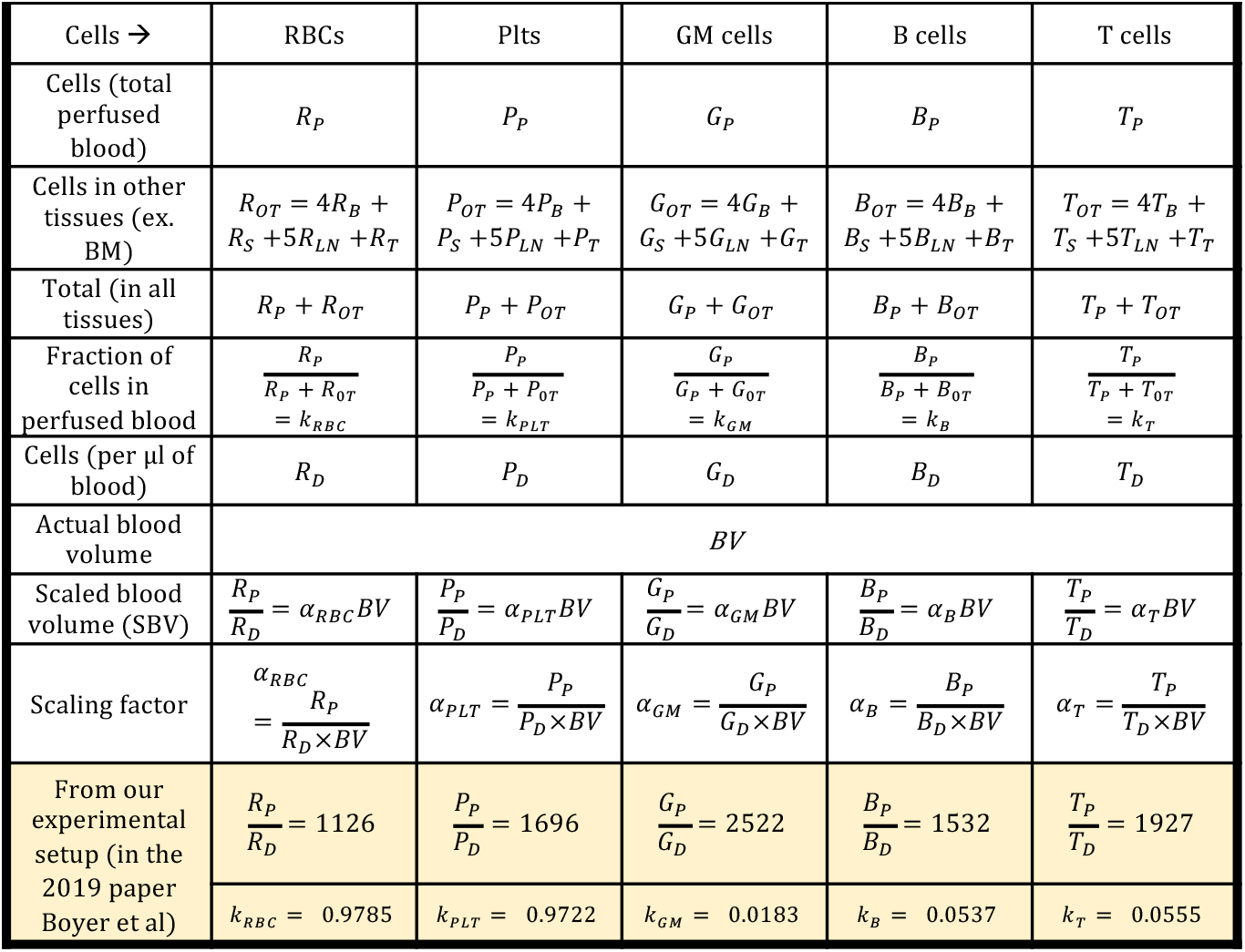

### BOX 3: Antibody-bead cocktail

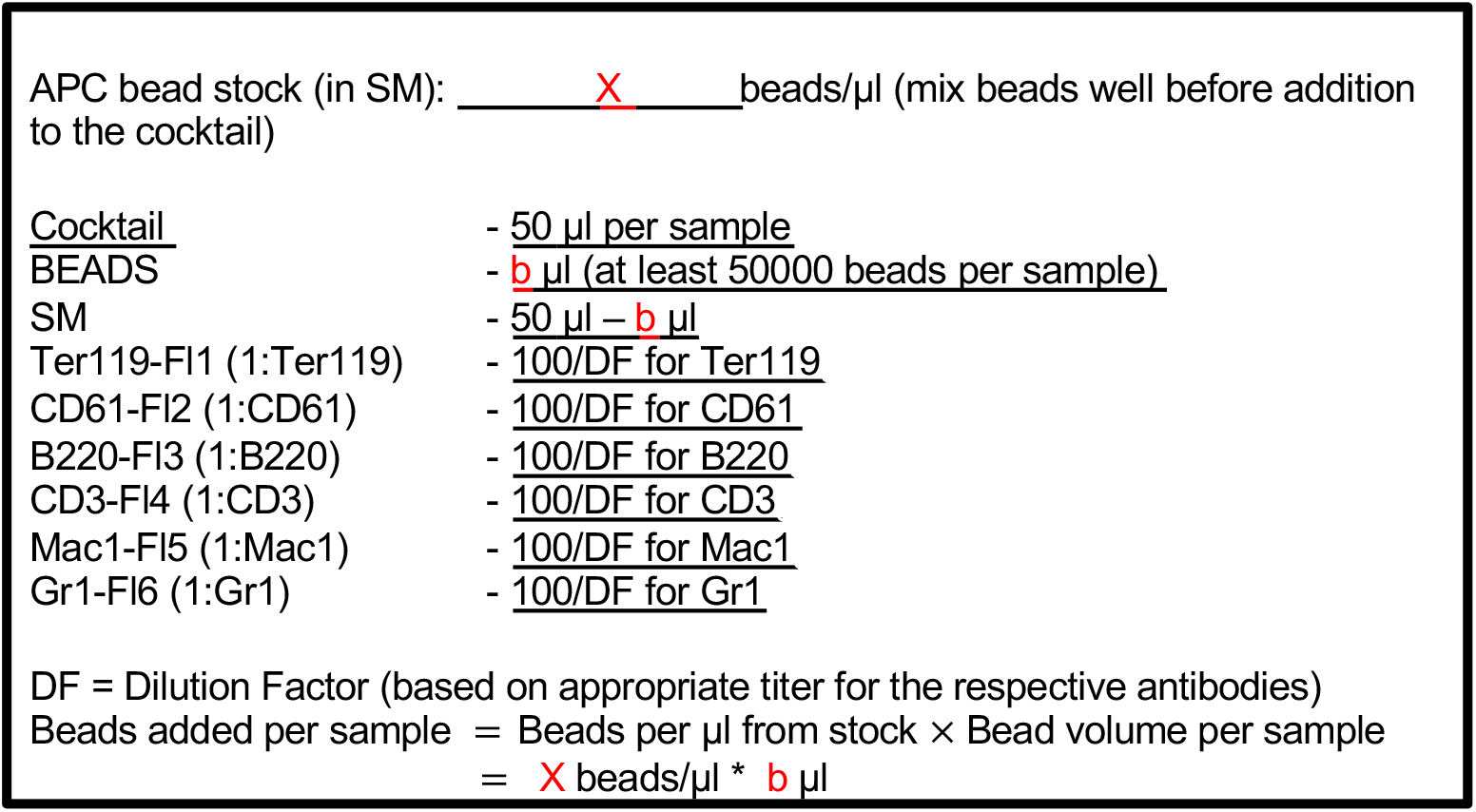

### BOX 4: Whole Blood Gating Strategy

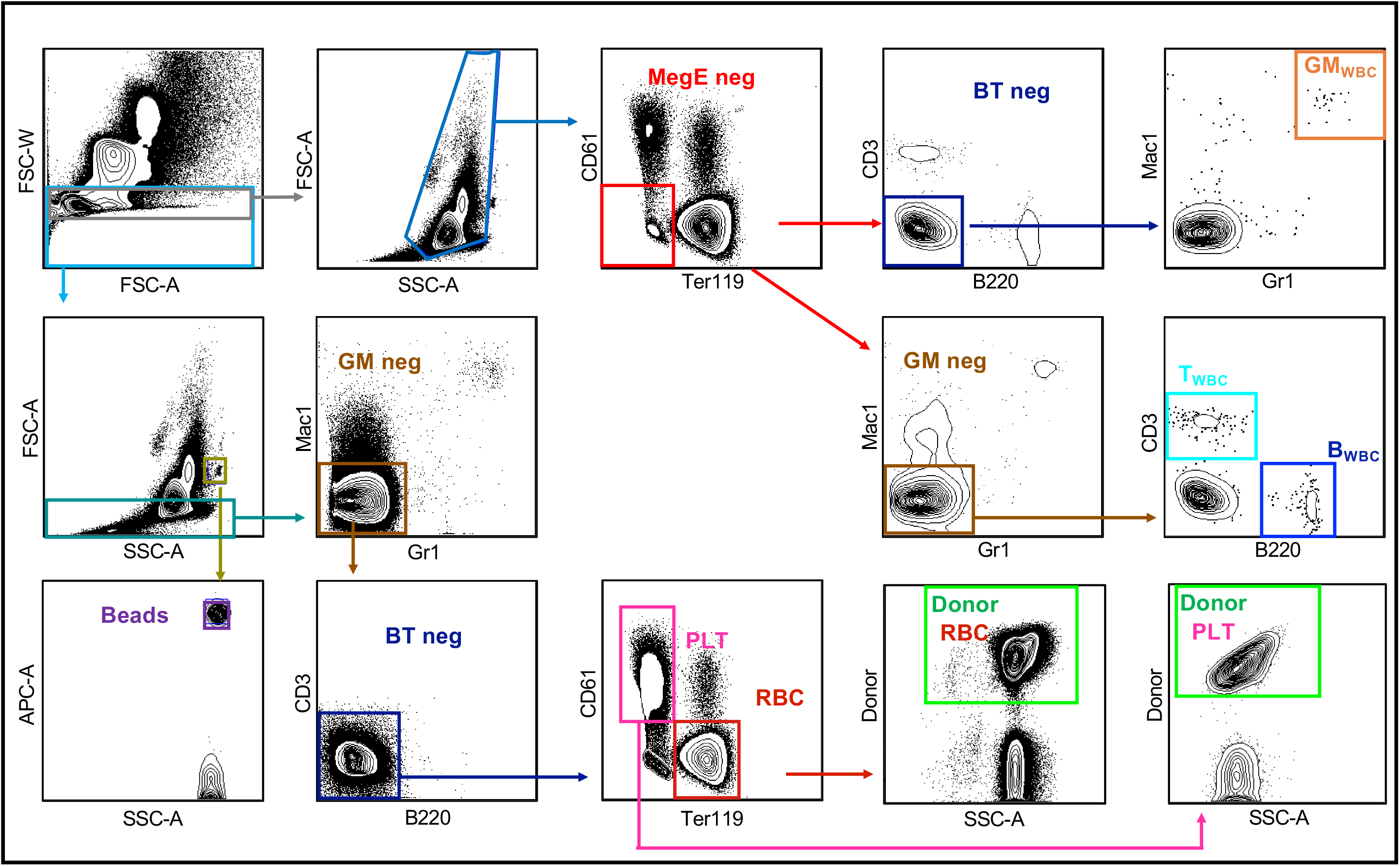

### BOX 5: ACK Lysed Blood Gating

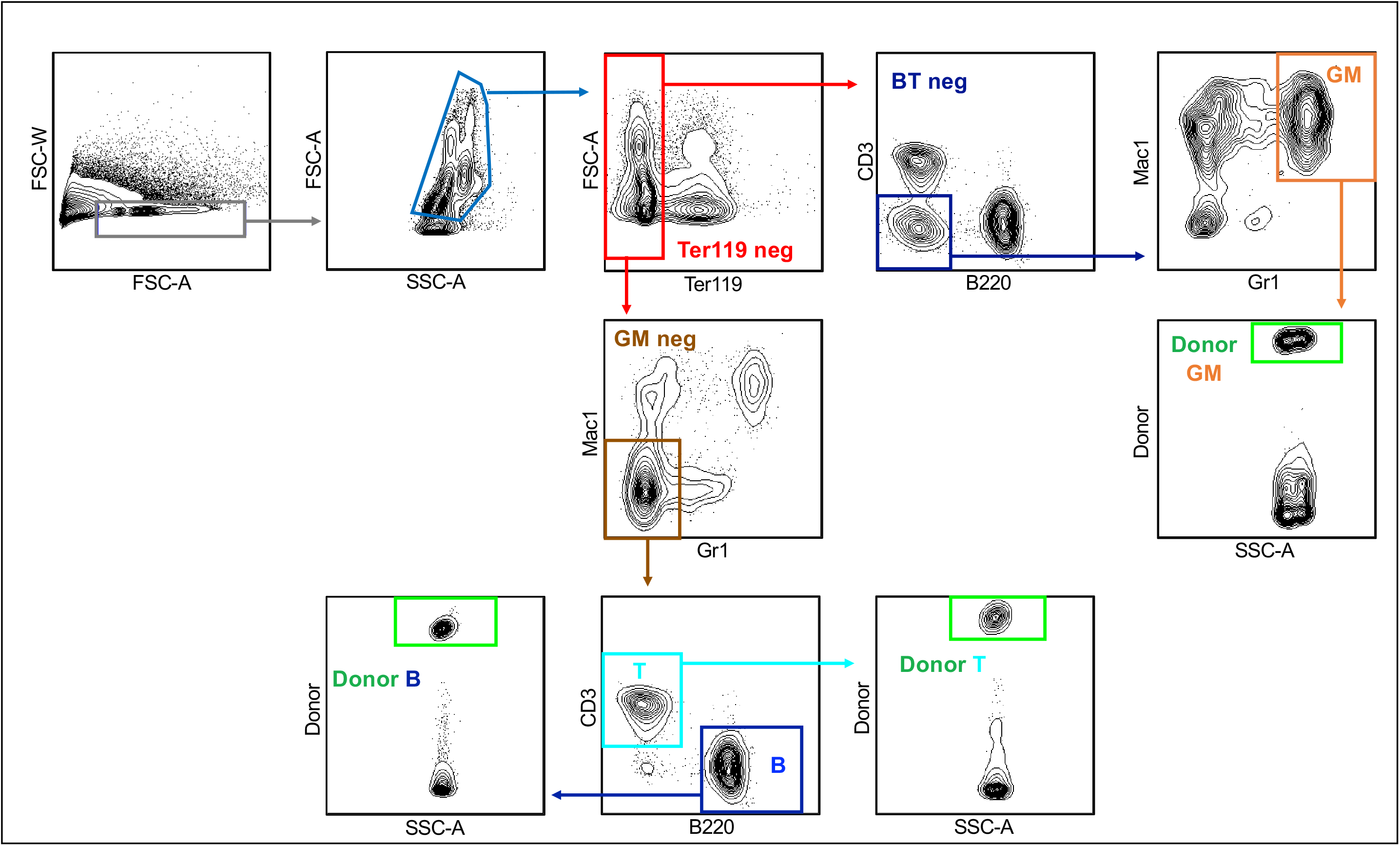

### BOX 6

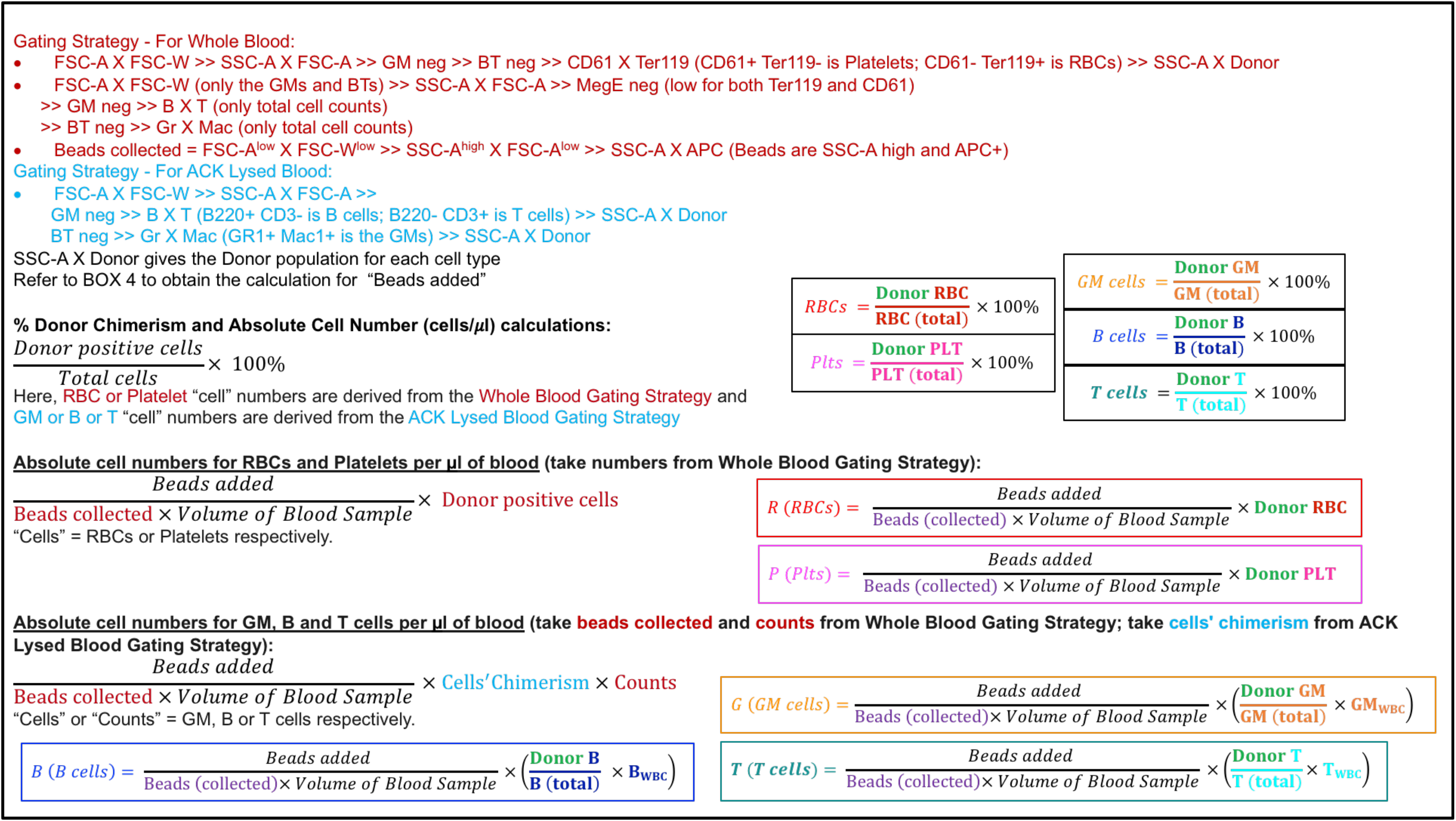

### BOX 7 Test: Conversion of cell numbers from tail bleed to “per mouse” cell number

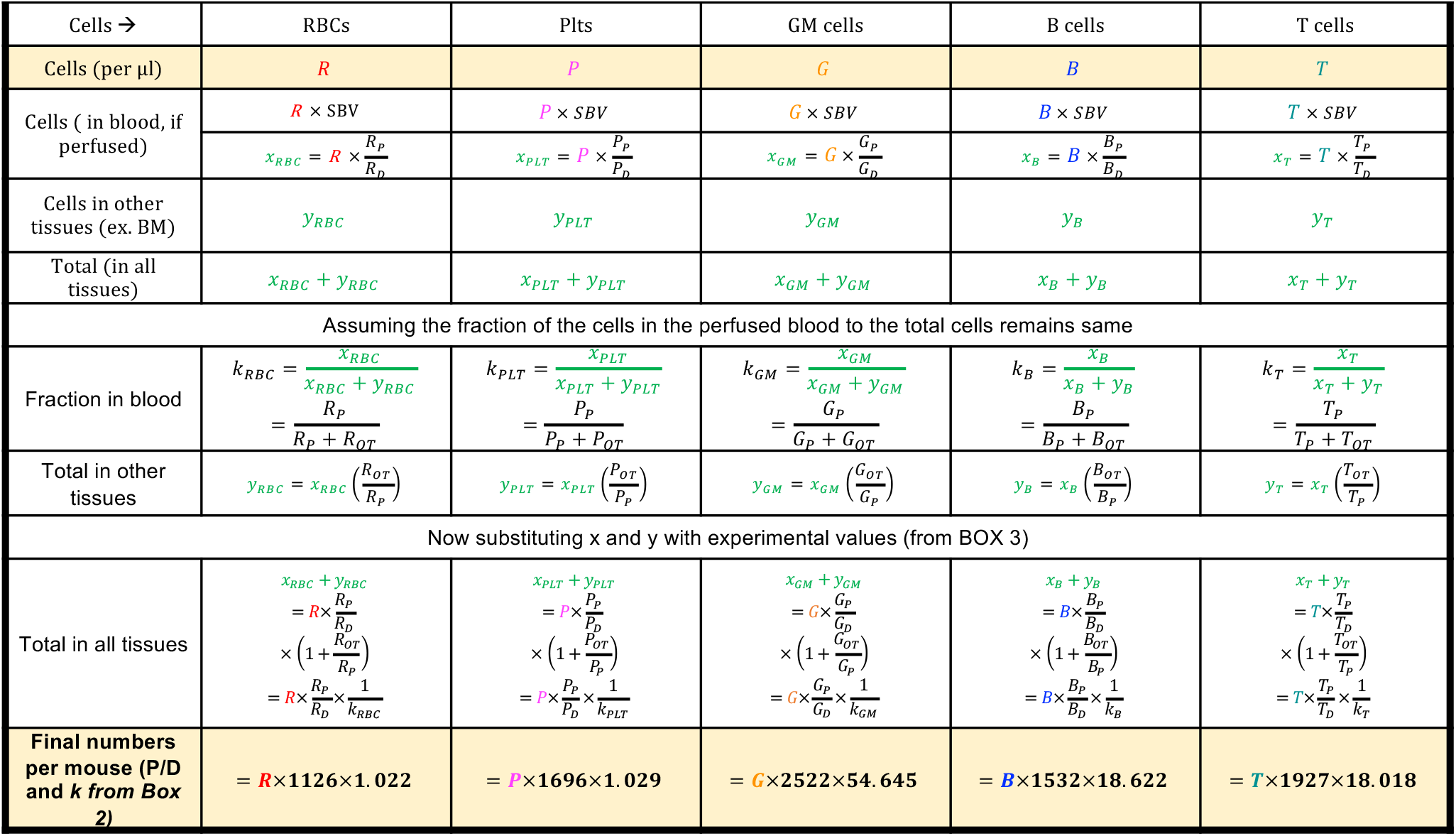

### BOX 8

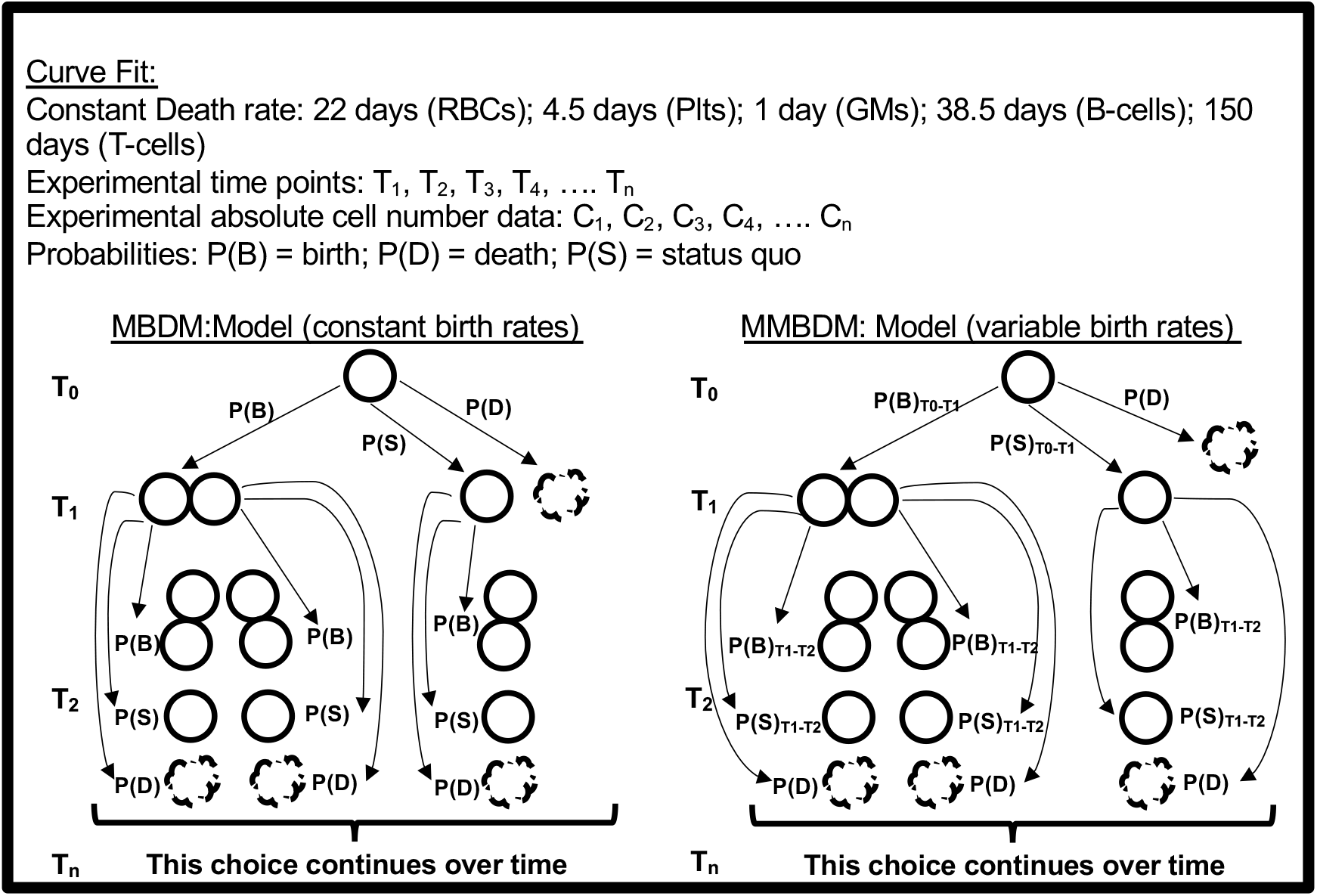

